# Biologically informed neural network models are robust to spurious interactions via self-pruning

**DOI:** 10.1101/2025.10.24.684155

**Authors:** Olof Nordenstorm, Hratch Baghdassarian, Xuechun Xu, Douglas A Lauffenburger, Avlant Nilsson

## Abstract

Computational models of cellular networks hold promise to uncover disease mechanisms and guide therapeutic strategies. Biology-informed neural networks (BINNs) is an emerging approach to create such models by combining the predictive power of deep learning with prior knowledge, a vital aspect of biological research. The architectures of BINN’s enforces a network structure from which mechanism can ideally be inferred. However, a key challenge is to evaluate the reliability of these models, as cells are inherently complex, involving intricate and sometimes unknown interactions. Currently, analysis mainly focuses on selected pathways rather than a more comprehensive perspective. In this work we demonstrate an alternative holistic approach: we measure to which extent purposefully introduced spurious interactions are down-weighted by a BINN during training (self-pruning). The metric suggested RRA (Relative Residual Area) allows for direct distribution comparison with perfect self-pruning achieved at zero and a failure to self-prune if above one. To enable rapid testing, we updated LEMBAS (Large-scale knowledge-EMBedded Artificial Signaling-networks), our recurrent neural network framework for intracellular signaling dynamics, with full GPU acceleration. Our implementation achieves a >7-fold speedup compared to the original while preserving predictive accuracy. We evaluated self-pruning in 3 different datasets and found that when spurious interactions are introduced at random, the model prunes these to a larger extent than those from the prior knowledge network (PKN), provided the model is regularized with a sufficiently large L2 norm. This suggests that BINNs can be robust to uncertainty in the PKN.

**Implementation and application:** Our implementation of LEMBAS is freely available under a MIT license at https://github.com/AvlantNilssonLab/LEMBAS_GPU. The models and results to generate the figures can be downloaded through https://zenodo.org/records/17425598.

## Introduction

Intracellular signaling pathways allow cells to process and react to their environment. This includes transmission of signaling events from ligand-receptor interactions to transcription factors (TFs)(1). Disruptions in signaling networks are common in diseases such as cancer(2–4), and therefore, understanding the systemic effects of such perturbations is central to deciphering disease mechanisms and developing new drugs(5). However, this is challenging for several reasons, including the vast combinatorial space of possible interaction in a signaling network, the non-linearity of these interactions, and inter-pathway signaling interactions across the network(6). Therefore, computational modeling approaches have high utility for understanding signaling activity and the potential to transform therapeutic approaches by enabling rapid evaluation of interventions(5,7).

Traditionally, modeling approaches, such as ordinary differential equations (ODEs) and Boolean models, have been used to simulate cellular signaling(8). These models are constructed using known molecular interactions in cells and thus serve not only as predictive tools but also have potential to reveal mechanistic relations. However, such models face limitations in scalability and expressiveness(9–12). Specifically, ODE methods require detailed and comprehensive knowledge of the parameters describing molecular interaction; and when this is not available, these parameters are highly computationally expensive to infer(13). Additionally, boolean models require extensive manual curation and are qualitative rather than quantitative by construction. As an alternative, artificial neural networks (ANNs) have emerged as a versatile and effective tool for approximating complex functions, with many biological applications(14).

Recently, ANNs that make use of prior biological knowledge have been developed. The aim of these models is to enable predictions that are (at least in principle) biologically interpretable and thereby bridge the gap between predictive accuracy and mechanistic insight. These models are termed Biologically Informed Neural Networks (BINNs)(5,15), analogous to Physics-Informed Neural Networks (PINNs). Whereas PINNs embed physical knowledge (e.g., differential equations) into their loss functions or model design to ensure physical realism in outputs(16), BINNs integrate biological priors such as interaction graphs into the architecture(15). For example, we developed a BINN termed **LEMBAS** (Large-scale knowledge-EMBedded Artificial Signaling-networks) that uses a recurrent neural network to simulate intracellular signaling and explain steady-state TF activities in response to ligands and other perturbations(17). It embeds known signaling interactions as structural constraints, mapping weights to real molecular interactions and setting all others to zero, making predictions interpretable and traceable(17,19). Unlike conventional deep learning models with randomly initialized parameters with no correspondence to biological processes, LEMBAS links predictions to molecular mechanisms, offering a path towards constructing interpretable models of disease such as cancer (5). And in general, BINNs has been proven to offer mechanistic insight in many biological tasks(17–20), suggesting that prior knowledge-informed networks could form a foundation for system biology analysis(17).

While BINNs hold promise to explain and predict biological data, several challenges that are associated with Graph Neural Networks (GNN) also translate to BINNs. One issue is misannotation of edges, which may be particularly challenging for biological systems where there are many methods to infer prior knowledge networks, and where the results can differ markedly, including spurious connections (false positives) as well as missing interactions (false negatives) (21). Additionally, cell types differ in the structure of their connectivity due differences in which pathways they express, annotation of this is limited, thus unrealistic interaction can emerge when using PKN that don’t separate cell types. Failure to use the correct graph structure may cause incorrect relations to emerge in the network. We reason that spurious connections may be less detrimental, as GNN models can (at least in principle) down-weight (prune) interaction but most implementations cannot introduce new edges(22,23), which also applies to BINNs(17,18). However, the propensity of these networks to down weight incorrect edges through training on experimental data is typically not systematically evaluated.

So far analysis of BINNs has mostly centered on its predictions for specific pathways (17,19,20,24,25), rather than on system level network properties. But analysis of the internal structure of deep learning networks can be a useful tool to understand how the algorithms behave during training(26,27). An example of this is *Grokking*(28), a phenomenon where after training a model for many epochs following an overfit to the training data, there is rapid progress to a perfect generalization on the test set. This is hypothesized to be the result of the model finding an efficient algorithmic-like solution to fit the data(29,30), and it has been observed that this is accompanied by a large fraction of the weights approaching zero, suggesting that the algorithmic solution constitutes a sparse network of strong interactions. We reason that this resembles the structured sparsity observed in biological networks and that therefore systems level analysis from the grokking literature may translate to the domain of BINNs.

Here we propose a systems-level, heuristic metric for network soundness that tests if false edges introduced in the network are suppressed during training, thus mimicking false positives in prior knowledge. We argue that in practice, performing well on this test is a necessary, but insufficient, requirement for system-level trust in PKN-based models. To systematically analyze how a BINN performs on this metric, we implemented a GPU-accelerated version of LEMBAS, that preserves accuracy while reducing training time. We find that false edges were consistently assigned negligible weights and were therefore effectively pruned over the course of training across synthetic and experimental datasets and across hyperparameter settings. While the work is inspired by grokking, where models shift from memorization to algorithmic solutions, we note that perfect recovery is unlikely in current BINN formulations, due to their extreme under parameterization. Additionally, there are structural differences between the formulations that may influence how training progresses. Indeed, we find that unlike the sharp transitions seen in grokking, pruning in this setting unfolded gradually. Overall, these results suggest BINNs are resilient to false interactions, supporting permissive prior inclusion and demonstrating their potential to infer mechanistic network structure.

## Results

### The reimplemented framework improves speed and preserves performance

Extensive training and sampling is needed to test if a BINN framework is robust to false positives in the prior knowledge network. To enable this we reimplemented our BINN framework LEMBAS (**Figure 1a**) with GPUs capabilities, expecting a marked speed up. It is important to note that the purpose of this implementation is not to reproduce a model with the exact same properties as the CPU version, rather produce a model that attains similar results but with a speed up from the GPU. In conjunction, we also made modifications to align it with more conventional machine learning approaches to regularization terms in the loss function, as well as streamlining the overall structure of the code base **(see Methods for details)**. After the reimplementation we found a perfect correlation in outputs and gradients when only training with MSE **(Supplementary Figure S1)**, confirming that the core computations remained unchanged. However, we also updated the terms that enforced; steady state, state diversity, and our method to prevent confinement to local minima during training (**Figure 1b**). These changes are expected to cause discrepancies between the original CPU-based version of LEMBAS and the new, GPU-enabled version based on pytorch using CUDA(31), and we therefore compared predictive performance along with computational speed of each implementation on two datasets that we previously used for training and benchmarking(17). Both of these datasets measure the transcriptional response to ligand stimulation in macrophages, one with low coverage (23 unique stimulation conditions) (32) and one with high coverage (systematic stimulation using 59 different ligands, with and without co-stimulation with lipopolysaccharide) (17). Models trained on these datasets were tested using cross validation (leave one out, and leave three out), using the same folds as in the original publication. A dense matrix multiplication was used as opposed to the old sparse multiplication used in the signaling state. This is due to sparse matrix multiplications in GPU only benefit at higher level of sparsity that what we have in our networks that is >99% sparsity (33).

**Figure 1.**
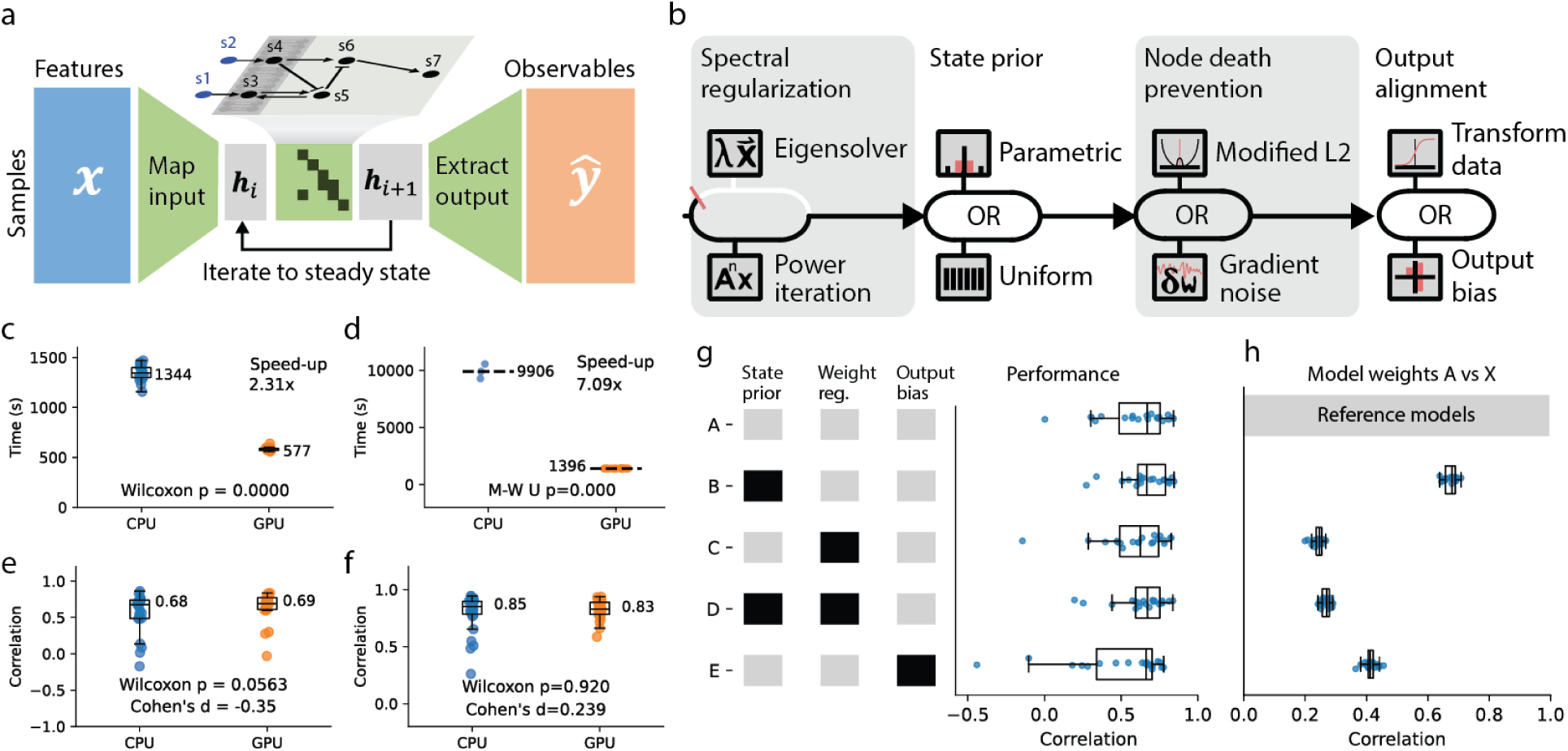
Reimplemented BINN framework with improved speed and preserved performance. **a)** LEMBAS is designed to propagate a hidden state that mirrors the dynamics of signaling molecules following real signaling events. The input to the model is mapped into the hidden state and the output is extracted from the hidden states. **b)** The modifications to LEMBAS compatible with GPU include spectral, state, and network weight (i.e., node death prevention) regularization. The new architecture is flexible with regularization term changes. For example, the state prior and node death prevention regularizations can be used in both a GPU and CPU implementation; however, the method to calculate the spectral radius is specific to GPU or CPU, using power iteration and an eigen solver, respectively. **c)** Comparison of runtime between CPU and GPU implementation of LEMBAS on the low coverage dataset(N=23), using the Willcox test. The black line represents the mean for each of the data sets. **d)** a similar comparison for the high coverage data set using the Mann-Whitney U-test on the CPU (N=3) and GPU (N=28) model. **e)** Comparison of performance on leaving one in the low-coverage data set (N=23). **f)** Comparison of predictive performance using cross validation high-coverage data set with 28 folds. **g)** Comparison of the original hyper parameterization to combinations of new modules, a black box indicates the new version **h)** Comparing the different conditions to the reference model A. Each model weights is compared to a model having A conditions with the same random seed.

For the low-coverage dataset, we preserved the hyperparameters from the CPU implementation. Here, runtime was reduced (*p* < 0.001, Wilcoxon test) to 42% from ∼23 min to 9 min (**Figure 1c**), on a high end desktop computer (see hardware). However, we expected that a switch to the new implementation would affect the optimal hyperparameters for both predictive performance and speed of training, as regularization is modified and GPUs can handle large batches more efficiently, owing to the inherent capacity for parallelization. We therefore manually tuned the hyperparameters to accommodate a larger batch in the high-coverage dataset, this was done by increasing the batch size and adjusting additional hyperparameter to get similar performance on the test set. This resulted in 7.1x faster training compared to the original implementation (*p* < 0.001, Mann-Whitney U-tes) from around 3 hours to 20 minutes (**Figure 1d**). Note that due to the long CPU runtime, we restricted the evaluation of runtime to three samples on the CPU and employed the Mann-Whitney U-test, which allows for unbalanced data sets. We reason that both analyses are relevant comparisons, but that the latter is more representative of the true speed gain, as using hyperparameters that were chosen to optimize performance on a distinct, old implementation with different regularization terms is not truly representative of how anyone would reasonably use a new implementation.

We found that the predictive performance was consistent between the CPU and GPU version. In the low-coverage setting **(Figure 1f)**, the GPU implementation yielded a median Pearson correlation of 0.69 among the folds compared to 0.68 for the CPU implementation. Interestingly, this improvement was nearly significant (p-value of 0.056, Wilcoxon test), despite limited hyperparameter tuning. For the high-coverage dataset **(Figure 1e)**, the median performance was 0.83 for the GPU and 0.85 for the CPU implementation (n.s., Wilcoxon test p-value = 0.92), in line with our expectation that predictive performance will be maintained in the new GPU version. Notably, the GPU implementation improved the mean performance of the models: correlation increased from 0.55 to 0.64 (Cohen’s effect size 0.35) and from 0.79 to 0.82 (Cohen’s effect size 0.239) in the low- and high-coverage datasets respectively, potentially due to fewer outliers with poor performance. While the calculated effect size is small, we note that it is present in both datasets.

### The reimplemented framework preserves both output and gradients

To follow up the observed differences in predictions, we investigated how the changes introduced affected output and gradients. One of these changes was that the CPU-dependent eigensolver for calculating the spectral radius loss, was replaced by a power iteration–based method previously used for RNNs(34–37). We found that five iterations provided a near-perfect correlation (∼0.95) with the CPU-based solver **(Supplementary Figure S2.)**. We also introduced a more exact approximation of the for state regularization. This new implementation correlated perfectly (r=0.9996) with the original implementation but results in larger magnitudes (x1.483) given the same hyperparameters **(Supplementary Figure S3)**. We also observed in preliminary testing that steady state and state regularization were no longer necessary for stable training as they were in the original CPU implementation. While the exact reason for this remains unclear, it may potentially be due to changes incurred from the new eigenvalue calculations. This prompted us to explore how removing these would affect predictive performance and runtime on the low-coverage ligand dataset. While the model could be trained faster (without regularization 523 seconds with 577 seconds and a p value of p<0.0001) **(Supplementary Figure S4)**, predictive performance did not significantly decrease (p = 0.445, Willcox test). **(Supplementary Figure S5)** This suggests that inclusion of these regularizations can be considered optional in this implementation.

Beyond assessing the direct effects of individual modifications, we also investigated their influence on learned parameters It is relevant to note here that the CPU model does not provide ground truth for what weights should be and that the parameters produced by the CPU and GPU models are likely equally useful from an interpretability perspective, rather we performed this analysis to highlight that some deviation emerge from the reimplementation.. We focused on three specific aspects: state regularization, weight regularization, and the inclusion of an additional bias on the output. The additional bias was, however, not used in the prior investigation, the bias has been introduced as an optional feature that can be included in the data and modelling demands it. These modifications represent true departures from the original implementation, in contrast to the power iteration method, which, if properly tuned, should reproduce the same eigenvalues as the original solver if so desired. No large departure in predictive accuracy was observed across all the conditions **(Figure 1g)**. While learned prior knowledge network weights remained largely unchanged following alterations to state regularization, adjustments to weight regularization induced a notable shift in the weight distributions. When both state and weight regularizations were modified simultaneously, deviations from the baseline were closest to the effect of weight regularization however it was slightly more pronounced. Notably, the effect of weight regularization appeared larger than that of introducing an additional output bias. While it could be expected that a modification to the model architecture would have a larger impact on parameters than a regularization term; the original L2 loss function was designed to strictly discourage zero weights and therefore its pronounced effects could be anticipated **(Figure 1h)**. A more detailed analysis comparing the weights of signaling **(Supplementary Figure S6)** and weights of the final output **(Supplementary Figure S7)**, shows that introduction of the output bias strongly affects the weights connecting the output of signaling to the model output, which is not surprising as when the output bias is introduced, the data is no longer sigmoid transformed as opposed to the other settings.

### False interactions vanish during training

False positives in the context of biological graphs can be viewed as the existence of entries connecting biomolecules that do not directly interact. It is well known that modern methods to produce these interaction graphs can produce inconsistent and incorrect results, this manifest as incorrect signs (activation/inhibition) in interactions (38), and false positives and false negatives in the adjacency graph(39). A neural network that is conditioned on such false interactions may potentially produce unrealistic structures that fail to model real biological processes. Ideally such false interactions should be removed during training, and it is standard practice in deep learning to simultaneously regularize weights (interactions) while optimizing the data fitting task. To investigate this self-pruning capacity of the framework, we added random edges to models and evaluate if these are pruned away during training (**Figure 2a**), the added edges where conditioned to conserve the connectivity distribution of the graph. For the sake of this analysis, we treat all stochastically added edges as false positives, even though theoretically the PKN may be missing interactions meaning that the added edges could be correct by accident in the real data setting. To test self-pruning in a setting where all added edges are certain to be false positives, we supplemented the 2 datasets previously described with an additional synthetic data set. It is generated using LEMBAS with the objective set to maximize signaling diversity within a network derived from KEGG(40), and therefore has a known, and theoretically recoverable parameterization. For the experimental datasets we expect the PKN to include many edges that are not expressed in macrophages, since the PKNs are compiled from observations among many different cell types. We will revisit this limitation in the discussion

**Figure 2.**
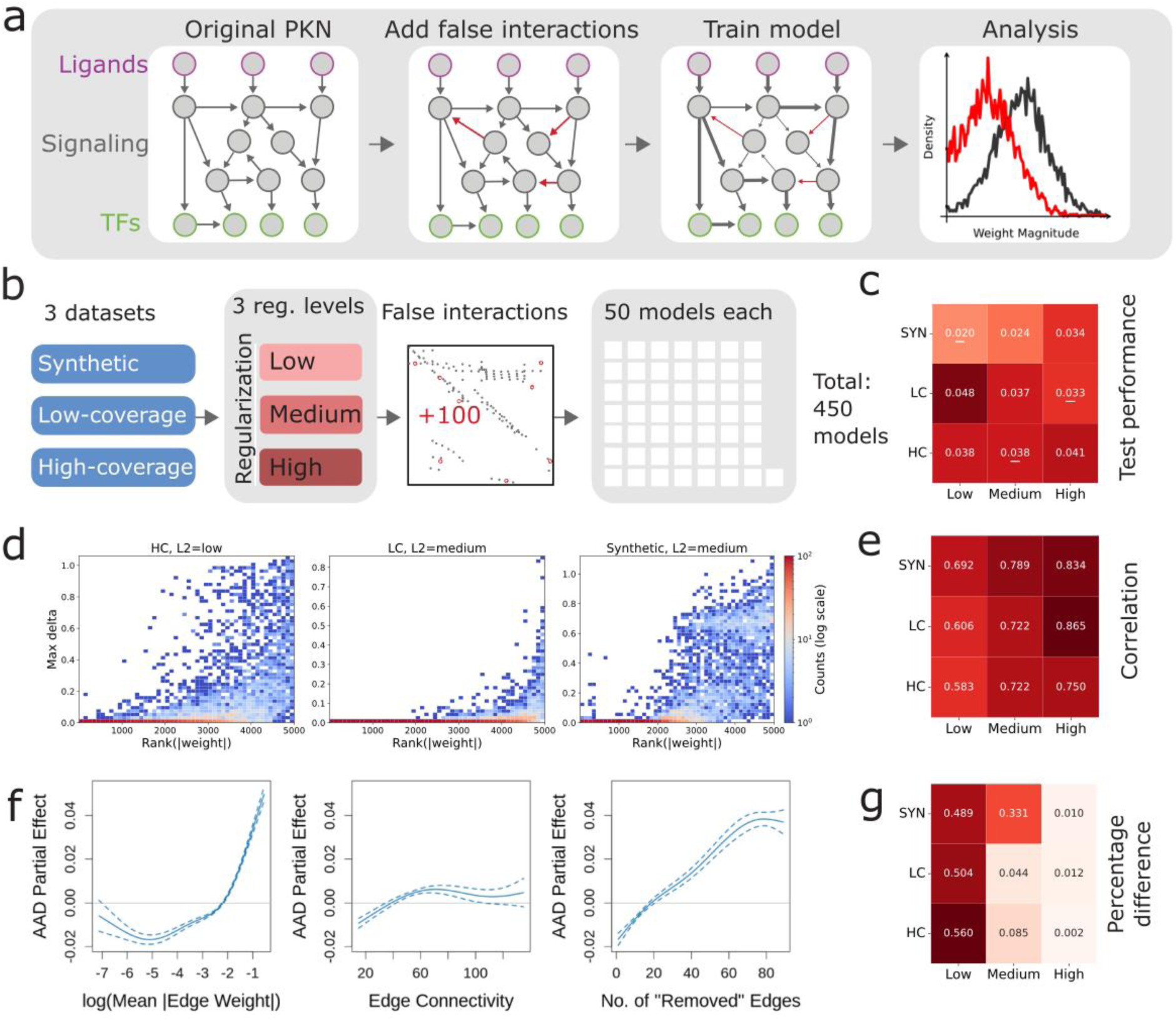
Effects of weight pruning. **a)** Proposed approach for testing the effects of false positives. **b)** For each dataset, three different levels of L2 regularization were tested, and for each combination, 50 models were trained with 100 false interactions each. **c)** The effect of regularization on test performance in the different datasets. A white line indicates the best result for each dataset. **d)** A heatmap of the effect of zeroing out a weight and the max delta effect. **e)** Correlation across all datasets between weight magnitude and the max delta effect, we find that there is generally high correlation with an increase as L2 is increased. **f)** Analysis of the generalized additive mixed model (GAMM) for the AAD indicates that the magnitude of the weights, together with the proportion of weights pruned to zero, exerts a greater influence on the model’s behavior than the network connectivity. Additionally zeroing a large weight can have an influence similar to zeroing 80 edges. **g)** Percentage difference as measured if all weights below the median is zeroed out.

For each dataset we investigated the effects of 3 different levels of L2 regularization (low (10E-6), medium (10E-5), and high (10E-4)). To gain statistical confidence, we performed this analysis in an ensemble of 50 models, with 100 stochastically added edges each (**Figure 2b**). For all datasets the uniform regulation of state was removed, as its main purpose is to improve gradient flow of the model, not to enforce known priors about the signaling process, and therefore enforcing it could promote false correlation between nodes to satisfy this objective. It is known that a too low regularization level permits overfitting, while a too high-level leads to underfitting. We therefore evaluated the test performance under the different regularization settings and found that the optimal level is dataset dependent (**Figure 2c, Supplementary Figure 8**). For the synthetic dataset, where most edges are expected to contribute to the solution, low regularization worked best, whilst high regularization worked best for the low coverage dataset, where only a subset of the network is expected to be activated. The high-coverage dataset achieved its optimal predictive performance in the medium regularization level; however, this performance was similar to the low level.

To assess the self-pruning ability in these settings, we examined how low-magnitude weights contributed to the models ‘predictions. Our rationale for this was: if small weights meaningfully affect the output, then pruning based on weight norms would be misguided, as the magnitudes would not imply how meaningful an edge is. To test this, we zeroed out induvial weights and observed the max delta of this model compared to the non-disputed model on the entire data set, overall small weights did not seem to have a large effect **(Figure 2d)**. Additionally, we tested the correlation between the weight zeroed out and max delta, we see high correlation and increasing correlation when L2 is increased **(Figure 2e),** indicating that weight is a good proxy for importance in this metric. Because weight magnitude alone does not guarantee mechanistic relevance, we tested whether and which properties of the PKN are associated with LEMBAS’ predictive performance, consistent with functional relevance. Specifically, we asked whether learned edge weights have a significant association with AAD (Absolute Average Deviation) independent of network topology, as represented by edge connectivity. To this end, we fit generalized additive mixed models (GAMMs) with AAD as the response, modeling non-linear effects of learned edge weight and edge connectivity while controlling for dataset, L2 regularization strength, number of edges removed, and edge type (stochastic vs true), and including model identity as a random effect **(Figure 2f).**

We found that, after accounting for network topology, edge weight magnitude shows a significant association (F-test p < 2.2e-16) with LEMBAS’ prediction sensitivity. We confirmed this using a likelihood ratio test (LRT) comparing the full model to a reduced model excluding edge weight. The LRT was highly significant (χ² p < 2.2e-16), indicating that edge weight captures additional explanatory information beyond network topology and increases the proportion of deviance explained by 5.62 percentage points. To assess whether these effects exceeded what would be expected by random chance, we compared the observed model F-statistic and LRT deviance to null distributions generated by permuting AAD 1000 times. Both statistics were highly significant under a one-sided permutation test (p < 1e-3).

Next, we investigate if a natural pruning point exists, i.e if all weights below a certain magnitude can be removed without detrimental effect to performance. However, no such point could be found that could be used across L2 conditions and datasets exemplified by the diversity observed at a median cutoff **(Figure 2g).**

### A metric for self-pruning in BINNs

To track performance across training, we saved the model at regular intervals (checkpoints), along with the initial parameterization before training started. We recorded the weights of true and false edges and calculated their magnitude (absolute value). By plotting the CDF (Cumulative Distribution Function) of the weights in the middle checkpoint we find a clear separation between added and true edges upon visual inspection **(Figure 3a, Supplementary Figure S9).** To quantify this separation, we calculate the residual area i.e the area of the upper left corner between the CDF and the point (0,1.0) a larger area of this type imply tendency towards larger weights, we compare the distribution of these areas for the true and added edges with a Wilcox test. With this metric, we find significant self-pruning for all datasets and L2 settings except the two lower L2 settings for the high-coverage dataset **(Figure 3b, Supplementary Figure S10).** We inspect this residual area metric across training time, and find that the distributions do not deviate at the start of training, and that they begin to deviate as training proceeds, indicating that it is the training dynamics that result in the separation of residual areas **(Figure 3c, Supplementary Figure S11).** Generally, we also see that the dataset and L2 norm affect the overall weight norm of weights in a trained model. Thus, to compare performance across models we need a metric that accounts for the model-specific distribution of weights.

**Figure 3.**
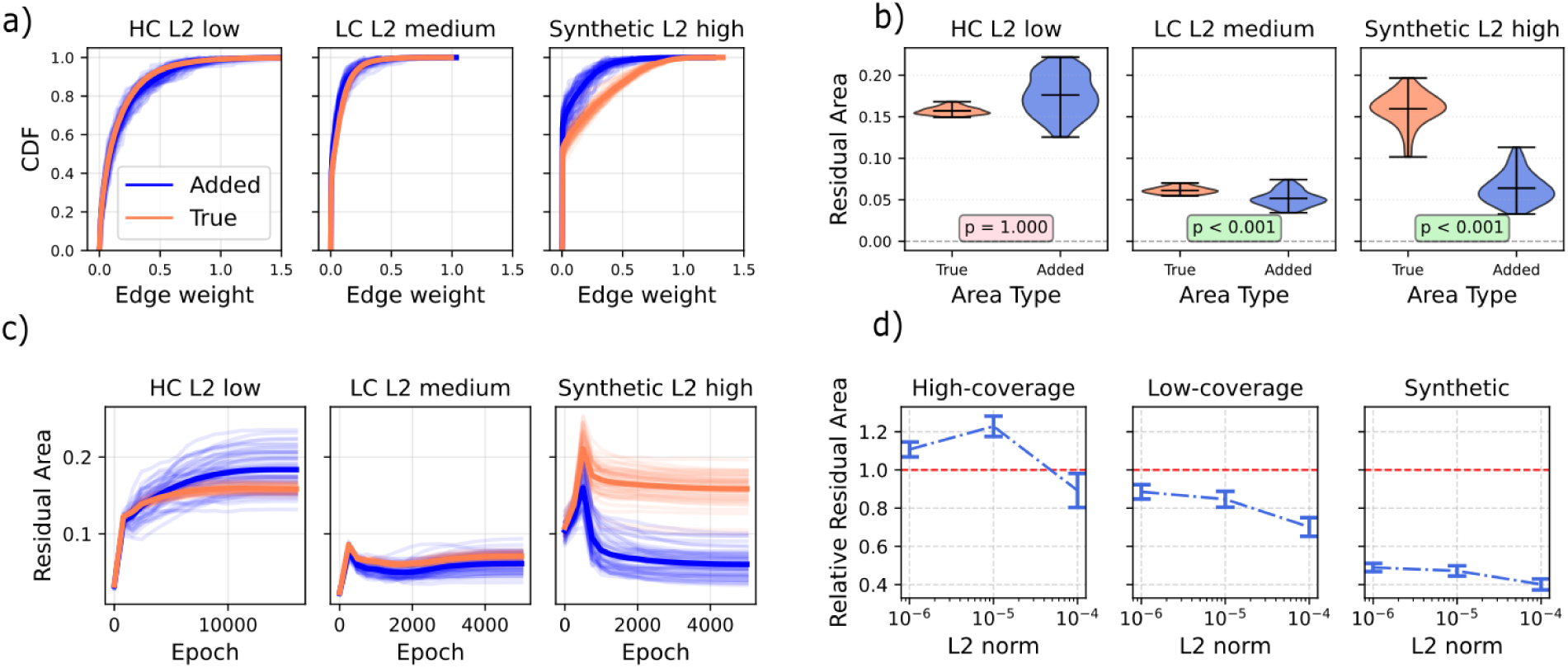
Self-pruning over models, time and L2 norm. **a)** The CDF of the weight magnitude of the added and true edges, across three datasets and three L2 norm. **b)** Violin plot of the residual area produced between the model across the same conditions. **c)** Residual area over training across the conditions. **d)** The relative residual area serve as metric for self-pruning we see that across all models we are experiencing some level of self-pruning with a sufficiently large L2 norm, we see a tendency that the larger the L2 norm the higher the self-pruning effect. The error bars indicate the significant of 0.05 and the red line is the self-pruning cutoff.

For this we introduce the relative residual area (RRA) to assess the self-pruning effect compared to the internal weight’s distribution of the model. For this we divide the residual area of added edges by the residual area of the true edges, introducing an internal baseline that allows comparison across conditions. This metric has clearly interpretable reference points, an RRA of zero implies perfect self-pruning, an RRA of 1 implies no self-pruning and an RRA of greater than one implies that added edges are preferred. Similar to our comparative analysis, using this metric, we find that a higher L2 tends to lead to a lower RRA and that in seven out of the nine tested conditions, we observe an RRA lower than one. The lowest RRA is in the synthetic dataset, which is expected as there is a perfect correspondence between the PKN and the relations expressed in the data, and the guarantee that the model is not under parameterized **(Figure 3d)**. Additionally, we made an analysis of self-pruning when varying the spectral radius **(Supplementary Figure S12-13)** and of including the output bias **(Supplementary Figure S14)**, observing limited effects on RRA,.

We also investigated if RRA can be used as a heuristic metric indicative of the mechanistic relevance of learned edge weights. For the synthetic setting where the ground truth is available, a direct measure can be constructed as the correlation between learned and ground truth weights. We found a correlation between the RRA score and this metric across models (r = –0.379, **Supplementary Figure S15**), indicating that lower RRA indeed generally reflects a higher similarity with ground truth. However, there was also considerable variability between these scores, indicating that further research into suitable mechanistic proxies could be needed. However, this is only possible to test synthetic models, as there is no ground truth available in the other setting. For these we have instead investigated if there is an association between a high weight and the number of references that support the edge in literature, we indeed find such an association **(Supplementary Figure S16),** this shows that more supported edges were more likely to be important in model prediction. These two analyses directly and indirectly support the RRA metric as a tool to assess mechanistic soundness.

## Discussion

In this work, we provide evidence that biologically informed neural networks (BINNs) can exhibit an emergent self-pruning behavior. For this we implement a new GPU version of our BINN framework LEMBAS and find that it is faster to train and produces comparable results. Additionally, the GPU implementation appears to generate fewer outliers with poor performance resulting in better mean performance. This re-implementation aligned the framework better with more recent methods in machine learning, including support for CUDA and new design choices such as the gradient noise, paving the way for future work on LEMBAS with an additional speed up. Together, these improvements enable a systematic investigation of self-pruning as an emergent property of the training of a BINN, and we test this by stochastically adding false-positive edges to the PKN and tracking their behavior during training. For this we used a synthetic dataset with a known ground truth, and on two datasets from stimulated macrophages. Across all datasets, we found that the distribution of weights learned for false edges changed over the course of training, and that self-pruning was enhanced by increased levels of L2 regularization.

Pruning and self-pruning is studied in the broader deep learning literature, particularly in the context of model compression(41) and as a theoretical tool to understanding how deep learning models learn(42). However, its emergence in the context of knowledge-informed network topologies has, to our knowledge, not been observed. This context introduces a key distinction: rather than pruning excess parameters in unconstrained architectures, here the network learns to denoise and refine biological prior knowledge, by actively down-weighting incorrect interactions in the knowledge network during training. Meaning that models set sufficiently small weights on edges as to effectively remove their effect, thus improving downstream interpretability work of the model as edges in the final model is a denoised PKN. When formulating this task, we were inspired by the results related to the grokking phenomenon. During Grokking, models trained on algorithmically simple tasks (e.g modular addition) (28) or on simple datasets (e.g classification on MNIST(43)) undergo a delayed phase transition: after an extended period of overfitting, the network abruptly discovers a generalizable, sparse representation that achieves near-perfect test accuracy. In these cases, it has been hypothesized (28) that the network synthesizes a compact algorithmic solution requiring only a small subset of weights, with the remainder becoming redundant and thus minimized. This hypothesis is supported by recent work (44) where the compact solution is particularly pronounced under L2 regularization, which also accelerates convergence by penalizing redundant parameters. In this work we observed that the training dynamics follow a more typical pattern: generalization improves steadily, and self-pruning appears to emerge as a by-product of regular training, potentially due to the sparsity already being enforced at the beginning of training. Nevertheless, it indicates that the redundant parameters introduced in addition to the true biological structure are being removed throughout training.

A key limitation in comparing BINNs to the systems normally studied in the Grokking literature stems from the intrinsic complexity of cells. Unlike synthetic tasks with compact solutions, cellular signaling is not expected to conform to the simplified functional forms of current BINNs such as LEMBAS. Consequently, despite its utility, self-pruning was not universal: in the high coverage dataset, the residual-area metric showed no preferential down-weighting of false edges at either of the two lower L2 regularization levels, despite the latter demonstrating the best predictive performance. Therefore self-pruning is more likely to be observed in stronger regularization regimes. Moreover, we reason that instances where self-pruning is not observed mat exemplify two main limitations: only a subset of all cellular processes are included in the models; and models naturally make simplifying assumptions that do not holistically capture all the intricacies of the biological mechanisms they represent. For example, in LEMBAS the contribution of each interaction is modeled independently, ignoring the interdependent relations between molecules. Another example is subcellular compartmentalization, which is ignored in LEMBAS but critical in real cells. As a result, the model is heavily under parameterized and LEMBAS instead uses the network edge weights to compensate for this lack of complexity. This is supported by the fact that, when training on synthetic data, the model shows stronger pruning, consistent with its well-parameterized functional structure. It is therefore encouraging that self-pruning emerges despite these constraints, suggesting that BINNs still learn meaningful approximations to underlying biological mechanisms. This points to their potential not only as predictive tools but also as engines for hypothesis testing and rejection in cellular systems. Here, the architectural structure is more important than the exact parameter count: a large, dense network without biological grounding, while technically not under parameterized, may fit data but would remain opaque and less mechanistically informative. So, to meaningfully mirror biological systems and learn mechanistically plausible representations, BINNs must strike a balance between expressiveness and structural constraints.

As highlighted in many recent articles (5,7,15), the field of BINNs is in need of robust evaluation frameworks and systematic strategies for hypothesis generation. We propose that the emergence of self-pruning holds promise of progress for both challenges. Specifically, the selective removal of non-contributory or biologically implausible edges during training can serve as an implicit validation mechanism. A framework that fails to prune added edges and instead relies on them for prediction, may indicate a failure to replicate biological reality, that would likely extend to false-positive interactions in the prior knowledge network (PKN). Frameworks that promote self-pruning enable a more permissive initial construction of PKNs that include uncertain interactions, with the expectation that training will refine this structure. This could be specifically useful in defining cell type specific interactomes as these interactomes are not fully mapped out and databases often have limited separation between cell types. Importantly, systems level metrics such as RRA proposed here provide a simple, architecture-agnostic test that does not rely on manual pathway curation, unlike many existing tests of biological feasibility. In LEMBAS, the RRA metric serves as a tool that partially assesses mechanistic relevance of learned edge weight magnitude. Consequently, the RRA can be modified and extended to use other metrics of edge-importance than weight magnitude, such as integrated gradients (45), where the distributions of the saliency between edges would serve as the feature compared, A limitation of the RRA metric is that it is defined relative to a reference model which could allow many real interactions to be pruned out as long as added edges are also pruned. We emphasize that self-pruning alone is limited in scope and should be combined with complementary metrics to assess how well models approximate the real mechanism inside of cells. We therefore interpret self-pruning as a system-level indication of biologically meaningful structure, rather than as direct evidence regarding individual edges.

## Methods

### Datasets

Throughout this paper we make use of three data sets: a high-coverage(17), low-coverage(32) and synthetic data set(17). The high and low coverage data set are both instances of macrophages simulated with ligands with a read out of gene expression. The synthetic data set was created using the CPU implementation of LEMBAS.

### Matrix multiplications and gradient calculations

We replaced the sparse matrix computations that were used in the original implementation of LEMBAS with dense matrix computations performed on a GPU, using python (v3.12.3) pytorch (v2.4.0), and cuda (v12.4). Originally, LEMBAS used accelerated sparse matrix computations that are possible on CPU, but the porting to a GPU required the use of dense operations. To ensure that only allowed interactions are maintained throughout training with the dense implementation, we expand our weight-update rule to set non-existing interaction to zero after each weights update has been performed. To make use of the sparse structure of the prior knowledge network, the original implementation of LEMBAS used a manually derived implementation of backpropagation and gradient calculation. The GPU-enabled version instead makes use of the standard autograd method that is integrated in pytorch.

### Spectral radius computation

Originally, a spectral radius loss term was introduced to ensure convergence to a steady state. For this, sparse eigenvalue calculations were performed using the scipy library, which is based on the ARnoldi PACKage (ARPACK) implementation(46). This method was not available on the GPU, and we replaced it with a power iteration method to approximate the dominant eigenvalue directly on the GPU. Instead of checking for convergence at each step of the algorithm, a fixed number of iterations (5) was used, as this reduces the computational overhead.

### Weight norm regularization and gradient noise

In this implementation, we penalize the model weights using standard L2 regularization. The original implementation included an additional term that also penalized weights near zero to prevent neuron death. In this implementation, neuron death is instead addressed by injection of gradient noise. We set the default regularization strength of this term to be the same as in the original CPU implementation (λ= 1e-6). To mitigate neuron death during training and potentially improve generalization, we applied gradient noise injection to parameters during training(47). The noise was drawn from a standard normal distribution scaled by a tunable hyperparameter. Formally, if *g* is the original gradient and ϵ∼N(0,I), the updated gradient becomes *g′=g+γɛ* where *γ* is the scaling hyperparameter. We set *γ* = 1e-9 for all analyses, except for cross validation on the hyperparameter optimized high-coverage ligand dataset, in which a varying noise scheme was used. Specifically, the noise was scaled according to the current learning rate: *g′=g+αγɛ* where *α* is the learning rate for this setup and we set *γ* = 1e-6. To measure the similarity of effect of the new regularization implementation on the steady state, a correlation was calculated on the gradient between the two implementations periodically (every 5 epochs) throughout training.

### State regularization

LEMBAS regularizes the predicted output to penalize deviation from a uniform distribution. This state regularization serves as a bayesian prior of diversity between samples. The original implementation penalized the mean and variance of a uniform distribution. The new implementation uses evenly spaced values (via linspace) to approximate a uniform distribution prior more directly. That is, we approximate the result of uniform distribution and penalize the deviation of the output from this distribution using MSE. We also penalize output values that fall outside the unit interval in alignment with the original implementation. A regularization strength of λ= 1e-6 was used for all analyses. To evaluate the similarity in the state regularization the gradients were stored and a final correlation was calculated at the end of the training. The methods was originally used in the CPU version of LEMBAS to enable faster training as without it the model had a hard time imposing significant variance in internal states however testing shows that this is no longer the case in the GPU version. The reason for this remains unclear but is likely due to the new gradient calculations.

### Addition of bias term on output

To enhance model flexibility and address potential data biases, an optional bias term was incorporated in the output layer of LEMBAS. This removes the non-negativity requirement on the model output, bypassing the need to apply additional transformation on inferred or measured TF activities, which may include negative values. As such, the predicted TF activity can be matched up directly with the data. This means that data using this term is no longer be transformed with a sigmoid.

### Other minor modifications

The gradient clipping term was changed to a norm gradient as it is a less biased scaling step. The frequency of convergence checks within the signaling network’s RNN propagation was reduced from being performed every iteration to every 10 iterations. This is due to the check requiring an expensive synchronization operation in the GPU setting.

### Performance Evaluation and Comparisons

We employed Cross-Validation (CV) to evaluate the effect of each algorithmic modification, independently, and together. Performance was assessed using Pearson correlation between predicted values and data. Execution time was assessed by the wall time for each fold in the CV. For the low-coverage dataset, the original hyperparameters were used to isolate for effects caused by the GPU implementation directly. For the high-coverage dataset, hyperparameter optimization was manually performed to decrease computational time while retaining predictive performance. This focused mainly on gradient noise schedules and batch size as these were found to drive most of the variance in performance. For the high-coverage dataset the CPU version of LEMBAS was run 3 times and used for the time comparison, for CV the reported correlations from the original publication were used.

### GAMM

All GAMMs were fitted using the R package *mgcv* (v1.9-4) with the number of basis functions for smooth terms set to *k* = 5. Partial effects and F-test p-values were obtained from models fit using restricted maximum likelihood (REML), while likelihood ratio tests were performed using maximum likelihood (ML) to ensure valid comparison of nested models differing in smooth terms. To model prediction sensitivity (AAD), we fit a GAMM with AAD as the response, including smooth terms for learned edge weight (log-transformed mean absolute weight), edge connectivity, and the number of edges removed, fixed effects for dataset, regularization strength, and edge type, and a random intercept for model identity. To assess whether the relationship between edge weight and AAD differed by edge type, we fit a second GAMM allowing the smooth effect of learned edge weight to vary by edge type, while retaining the same covariates and random-effects structure as the main model.43

### Self-pruning

To assess the sensitivity of the signaling module to weight pruning. We investigate how big of an effect removing all weights lower than a specific cutoff has on the final output.

We trained an ensemble of models for each dataset and level of L2 regularization, adding 100 stochastic edges to each PKN, in a conditioned way to preserve the connectivity distribution **(Supplementary Algorithm A1)**. The model weights were saved at checkpoints throughout training. The CDFs of edge weights were retrieved at the middle checkpoint. The hyperparameters for the model were the same as for the cross validation study. For the synthetic data set we reused the low-coverage models pick of hyperparameters, with state regularization removed.

### Hardware for simulations

Walltime comparison and CV on the low-coverage and high-coverage ligand dataset were performed on a Dell Precision 3680 with Intel(R) core(^TM^) i7-14700 cpu and an RTX 4000 Ada Generation Graphics Card. For the generation of self-pruning and cross validation on the high-coverage ligand dataset we used the Berzelius AI/ML cluster using several different GPUs. We also used Satori a computational resource supplied by MIT. Satori is a GPU dense, high-performance Power 9 system developed as a collaboration between MIT and IBM. It has 64 1TB memory Power 9 nodes. Each node hosts four NVidia V100 32GB memory GPU cards. Within a node GPUs are linked by an NVLink2 network that supports nearly 200GB/s bi-directional transfer between GPUs. A 100Gb/s Infiniband network with microsecond user space latency connects the cluster nodes together.

## Availability and Implementation

Our implementation of LEMBAS is freely available under a MIT license at https://github.com/AvlantNilssonLab/LEMBAS_GPU. The models and results to generate the figures can be downloaded through https://zenodo.org/records/17425598.

## Acknowledgements

We acknowledge funding from the SciLifeLab & Wallenberg Data-Driven Life Science Program grant no. KAW 2020.0239 (O.N, A.N.). The work of H.M.B and D.A.L. was supported in part by NIH grants (IMPAcTB contract #75N93019C00071, U19-AI167899, U19-AI135995). H.M.B. was supported by the MIT-Novo Nordisk Artificial Intelligence Postdoctoral Fellows Program and a Cancer Research Institute Immuno-Informatics Postdoctoral Fellowship (CRI12812). The computations were enabled by the supercomputing resource Berzelius provided by the National Supercomputer Centre at Linköping University and the Knut and Alice Wallenberg foundation.

## References

1. Kholodenko BN. Cell-signalling dynamics in time and space. Nat Rev Mol Cell Biol. 2006 Mar;7(3):165–76.

2. Sever R, Brugge JS. Signal Transduction in Cancer. Cold Spring Harb Perspect Med. 2015 Apr 1;5(4):a006098–a006098.

3. Zhong L, Li Y, Xiong L, Wang W, Wu M, Yuan T, et al. Small molecules in targeted cancer therapy: advances, challenges, and future perspectives. Signal Transduct Target Ther. 2021 May 31;6(1):201.

4. Tape CJ, Ling S, Dimitriadi M, McMahon KM, Worboys JD, Leong HS, et al. Oncogenic KRAS Regulates Tumor Cell Signaling via Stromal Reciprocation. Cell. 2016 May 5;165(4):910–20.

5. Nilsson A, Meimetis N, Lauffenburger DA. Towards an interpretable deep learning model of cancer. Npj Precis Oncol. 2025 Feb 14;9(1):1–7.

6. Hughey JJ, Lee TK, Covert MW. Computational modeling of mammalian signaling networks. WIREs Syst Biol Med. 2010;2(2):194–209.

7. Bunne C, Roohani Y, Rosen Y, Gupta A, Zhang X, Roed M, et al. How to build the virtual cell with artificial intelligence: Priorities and opportunities. Cell. 2024 Dec;187(25):7045–63.

8. Samaga R, Klamt S. Modeling approaches for qualitative and semi-quantitative analysis of cellular signaling networks. Cell Commun Signal. 2013 June 26;11(1):43.

9. Kapfer EM, Stapor P, Hasenauer J. Challenges in the calibration of large-scale ordinary differential equation models. IFAC-Pap. 2019 Jan 1;52(26):58–64.

10. How to deal with parameters for whole-cell modelling [Internet]. [cited 2025 May 23]. Available from: https://royalsocietypublishing.org/doi/epdf/10.1098/rsif.2017.0237

11. Hyduke DR, Palsson BØ. Towards genome-scale signalling network reconstructions. Nat Rev Genet. 2010 Apr;11(4):297–307.

12. Invergo BM, Beltrao P. Reconstructing phosphorylation signalling networks from quantitative phosphoproteomic data. Essays Biochem. 2018 Aug 2;62(4):525–34.

13. Kreutz C. Guidelines for benchmarking of optimization-based approaches for fitting mathematical models. Genome Biol. 2019 Dec 16;20(1):281.

14. Sapoval N, Aghazadeh A, Nute MG, Antunes DA, Balaji A, Baraniuk R, et al. Current progress and open challenges for applying deep learning across the biosciences. Nat Commun. 2022 Apr 1;13(1):1728.

15. Selby DA, Sprang M, Ewald J, Vollmer SJ. Beyond the black box with biologically informed neural networks. Nat Rev Genet. 2025 Mar 4;1–2.

16. Cuomo S, Di Cola VS, Giampaolo F, Rozza G, Raissi M, Piccialli F. Scientific Machine Learning Through Physics–Informed Neural Networks: Where we are and What’s Next. J Sci Comput. 2022 July 26;92(3):88.

17. Nilsson A, Peters JM, Meimetis N, Bryson B, Lauffenburger DA. Artificial neural networks enable genome-scale simulations of intracellular signaling. Nat Commun. 2022 June 2;13(1):3069.

18. Elmarakeby HA, Hwang J, Arafeh R, Crowdis J, Gang S, Liu D, et al. Biologically informed deep neural network for prostate cancer discovery. Nature. 2021 Oct;598(7880):348–52.

19. Meimetis N, Lauffenburger DA, Nilsson A. Inference of drug off-target effects on cellular signaling using interactome-based deep learning. iScience. 2024 Apr;27(4):109509.

20. Interpreting biologically informed neural networks for enhanced proteomic biomarker discovery and pathway analysis | Nature Communications [Internet]. [cited 2025 Sept 15]. Available from: https://www.nature.com/articles/s41467-023-41146-4

21. Stock M, Losert C, Zambon M, Popp N, Lubatti G, Hörmanseder E, et al. Leveraging prior knowledge to infer gene regulatory networks from single-cell RNA-sequencing data. Mol Syst Biol. 2025 Mar 3;21(3):214–30.

22. Kipf TN, Welling M. Semi-Supervised Classification with Graph Convolutional Networks [Internet]. arXiv; 2017 [cited 2025 July 9]. Available from: http://arxiv.org/abs/1609.02907

23. Veličković P, Cucurull G, Casanova A, Romero A, Liò P, Bengio Y. Graph Attention Networks [Internet]. arXiv; 2018 [cited 2025 July 9]. Available from: http://arxiv.org/abs/1710.10903

24. Kim Y, Lee H. PINNet: a deep neural network with pathway prior knowledge for Alzheimer’s disease. Front Aging Neurosci. 2023 July 14;15:1126156.

25. Liu X, Tao Y, Cai Z, Bao P, Ma H, Li K, et al. Pathformer: a biological pathway informed transformer for disease diagnosis and prognosis using multi-omics data. Bioinformatics. 2024 May 1;40(5):btae316.

26. Frankle J, Carbin M. The Lottery Ticket Hypothesis: Finding Sparse, Trainable Neural Networks [Internet]. arXiv; 2019 [cited 2025 Sept 15]. Available from: http://arxiv.org/abs/1803.03635

27. Nakkiran P, Kaplun G, Bansal Y, Yang T, Barak B, Sutskever I. Deep Double Descent: Where Bigger Models and More Data Hurt [Internet]. arXiv; 2019 [cited 2025 Sept 15]. Available from: http://arxiv.org/abs/1912.02292

28. Power A, Burda Y, Edwards H, Babuschkin I, Misra V. Grokking: Generalization Beyond Overfitting on Small Algorithmic Datasets [Internet]. arXiv; 2022 [cited 2025 Mar 27]. Available from: http://arxiv.org/abs/2201.02177

29. Liu Z, Kitouni O, Nolte N, Michaud EJ, Tegmark M, Williams M. Towards Understanding Grokking: An Effective Theory of Representation Learning.

30. Nanda N, Chan L, Lieberum T, Smith J, Steinhardt J. Progress measures for grokking via mechanistic interpretability [Internet]. arXiv; 2023 [cited 2025 May 26]. Available from: http://arxiv.org/abs/2301.05217

31. Scalable Parallel Programming with CUDA.

32. Xue J, Schmidt SV, Sander J, Draffehn A, Krebs W, Quester I, et al. Transcriptome-Based Network Analysis Reveals a Spectrum Model of Human Macrophage Activation. Immunity. 2014 Feb 20;40(2):274–88.

33. Mishra A, Latorre JA, Pool J, Stosic D, Stosic D, Venkatesh G, et al. Accelerating Sparse Deep Neural Networks [Internet]. arXiv; 2021 [cited 2025 Aug 26]. Available from: http://arxiv.org/abs/2104.08378

34. Yoshida Y, Miyato T. Spectral Norm Regularization for Improving the Generalizability of Deep Learning [Internet]. arXiv; 2017 [cited 2025 Sept 15]. Available from: http://arxiv.org/abs/1705.10941

35. Sengupta B, Friston KJ. How Robust are Deep Neural Networks? [Internet]. arXiv; 2018 [cited 2025 Sept 15]. Available from: http://arxiv.org/abs/1804.11313

36. Rosen Y, Kirsch L, Louzoun Y. Optimal network modification for spectral radius dependent phase transitions. New J Phys. 2016 Sept;18(9):093039.

37. Mak KL, Peng JG, Xu ZB, Yiu KFC. A new stability criterion for discrete-time neural networks: Nonlinear spectral radius. Chaos Solitons Fractals. 2007 Jan 1;31(2):424–36.

38. Türei D, Korcsmáros T, Saez-Rodriguez J. OmniPath: guidelines and gateway for literature-curated signaling pathway resources. Nat Methods. 2016 Dec;13(12):966–7.

39. Mahdavi MA, Lin YH. False positive reduction in protein-protein interaction predictions using gene ontology annotations. BMC Bioinformatics. 2007 July 23;8:262.

40. Kanehisa M, Goto S. KEGG: kyoto encyclopedia of genes and genomes. Nucleic Acids Res. 2000 Jan 1;28(1):27–30.

41. Wang H, Ma S, Dong L, Huang S, Wang H, Ma L, et al. BitNet: Scaling 1-bit Transformers for Large Language Models [Internet]. arXiv; 2023 [cited 2025 Aug 11]. Available from: http://arxiv.org/abs/2310.11453

42. Frankle J, Carbin M. The Lottery Ticket Hypothesis: Finding Sparse, Trainable Neural Networks [Internet]. arXiv; 2019 [cited 2025 Aug 15]. Available from: http://arxiv.org/abs/1803.03635

43. Liu Z, Michaud EJ, Tegmark M. OMNIGROK: GROKKING BEYOND ALGORITHMIC DATA. 2023;

44. Liu Z, Kitouni O, Nolte N, Michaud EJ, Tegmark M, Williams M. Towards Understanding Grokking: An Effective Theory of Representation Learning.

45. Mukund Sundararajan, Ankur Taly, and Qiqi Yan. 2017. Axiomatic attribution for deep networks. In Proceedings of the 34th International Conference on Machine Learning - Volume 70 (ICML’17). JMLR.org, 3319–3328.

46. Virtanen P, Gommers R, Oliphant TE, Haberland M, Reddy T, Cournapeau D, et al. SciPy 1.0: fundamental algorithms for scientific computing in Python. Nat Methods. 2020 Mar;17(3):261–72.

47. Neelakantan A, Vilnis L, Le QV, Sutskever I, Kaiser L, Kurach K, et al. Adding Gradient Noise Improves Learning for Very Deep Networks [Internet]. arXiv; 2015 [cited 2025 Mar 28]. Available from: http://arxiv.org/abs/1511.06807

