## Supplemental material for "Biologically informed neural network models are robust to spurious interactions via self-pruning"

### False interactions vanish during training of a biologically informed neural network

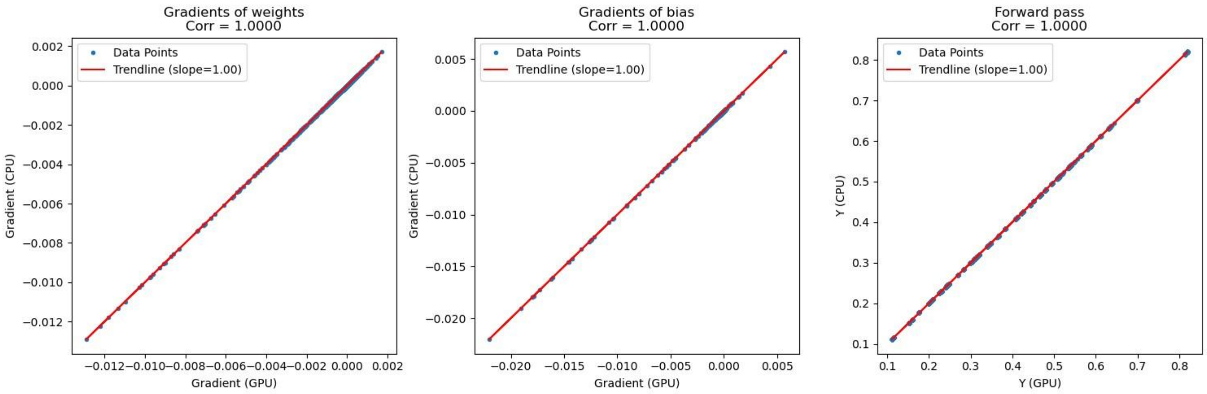


**Supplementary Figure S1. Comparison of CPU and GPU implementation on output**

A direct comparison of the CPU method and the GPU method with the same model parameters. For this we performed a comparative analysis of a single forward and backward propagation for each algorithmic modification under identical parameter settings. A model was trained on the GPU version of LEMBAS, and training was interrupted at an arbitrary intermediate step. The model's state was saved, and forward and backward propagations were executed with both the CPU and GPU-enabled version. The final results indicate perfect agreement between the methods with a correlation of one.

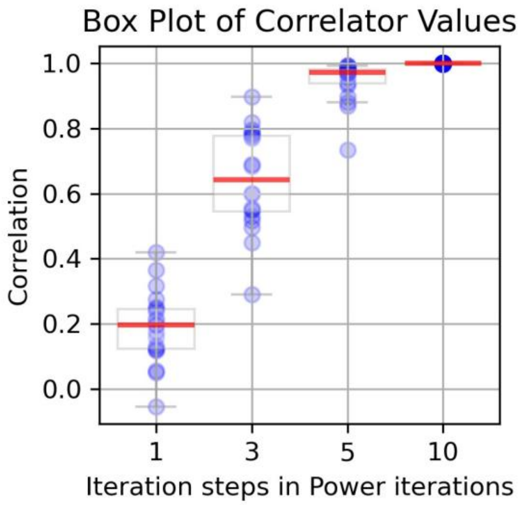


**Supplementary Figure S2. Evaluation of the power iteration-based eigensolver.** We needed to replace the eigensolver for over steady state regularization. This was necessitated due to a lack of GPU equivalent for the sparse eigensolver used in the GPU implementation (the ARPACK-NG-based eigensolver accessed through the SciPy package), which we replaced with a custom implementation of power iteration-based eigensolver. The power iteration algorithm is used to approximate the dominant eigenvector (associated with the largest eigenvalue in magnitude) of a matrix. It begins with a set of randomly initialized vectors. At each iteration, the algorithm multiplies these vectors by the matrix, effectively amplifying components in the direction of the dominant eigenvector. After each multiplication, the vectors are normalized to avoid numerical instability in case of divergence. By repeating this process, the vectors gradually align with the dominant eigenvector. Using multiple vectors in parallel helps accelerate convergence, since the initial ensemble better spans the vector space and increases the chance that some components already point to a large degree in the direction of the dominant eigenspace. Once the vector has approximately converged, the associated eigenvalue can be estimated by computing the Rayleigh quotient. To ensure that the method gave a similar result with this less exact eigensolver we backpropagate gradients using both methods and calculate correlation for many different steps during training. Since power iteration converges to the dominant eigenvector with increasing iterations, we conducted a parameter study to examine how the correlation between our results and the eigensolver solution evolves with iteration count. To test we checked how an exact solver compared to our power iteration method with different sets of iterations step we deemed that five iterations is enough to get near perfect correlation on the backpropagation using the method. As expected we saw monotonic growth of the median correlation as iteration increased. We observed high concordance in the gradients used for spectral radius loss calculation, with a median Pearson correlation of approximately 0.95 after just 5 iterations, and near-perfect correlation thereafter**.** As each extra power iteration increases the running time for the algorithm we decided that five power iteration steps offer a good trade off between exact eigenvalue calculation and compute efficiency.

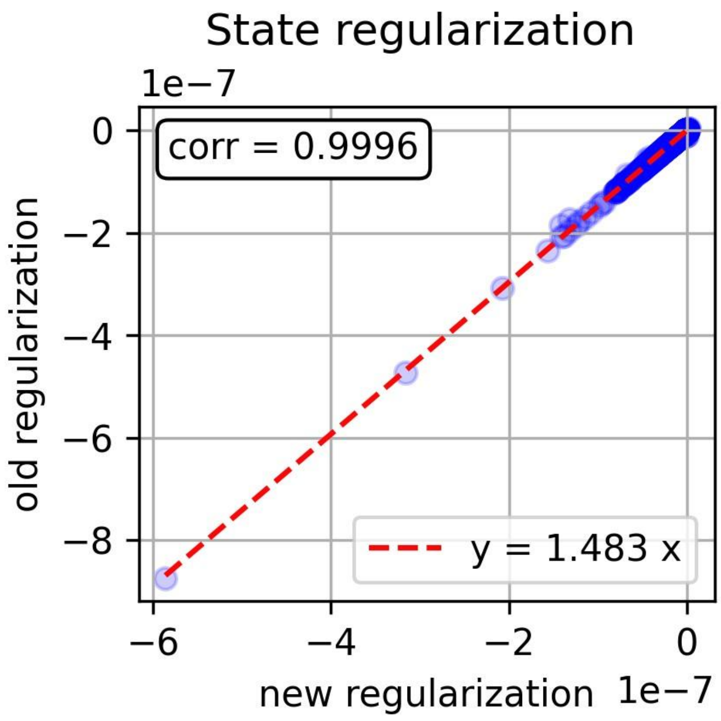


**Supplementary Figure S3. Comparison between state regularization methods.** In this figure we see the effect of the new state regularization. We see generally good correlation but note that there is a difference in magnitude between the gradients. This new implementation is a more strict approximation of a uniform distribution instead of the former method that only penalized on the mean and standard deviation of a uniform distribution. We compared our two versions of the state loss, the Pearson correlation of gradients between the state regularization methods where near ~~zero~~ one however there is a change in overall magnitude that affected the gradient of the model.

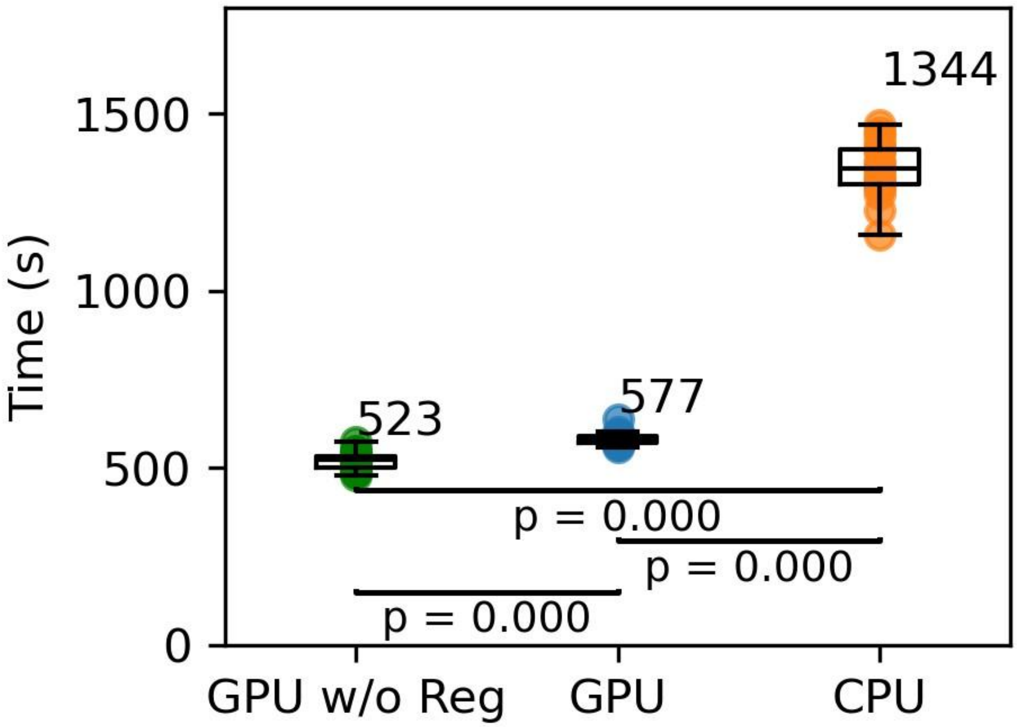


**Supplementary Figure S4. Effect of removing regularization on wall time.** After removing the state and steady state regularization terms we did see an increase in training time. This effect was significant however because of the minor decrease observed **(sup figure 5)**. We decided to keep the loss term in the algorithm even though it had a small effect on the speed performance.

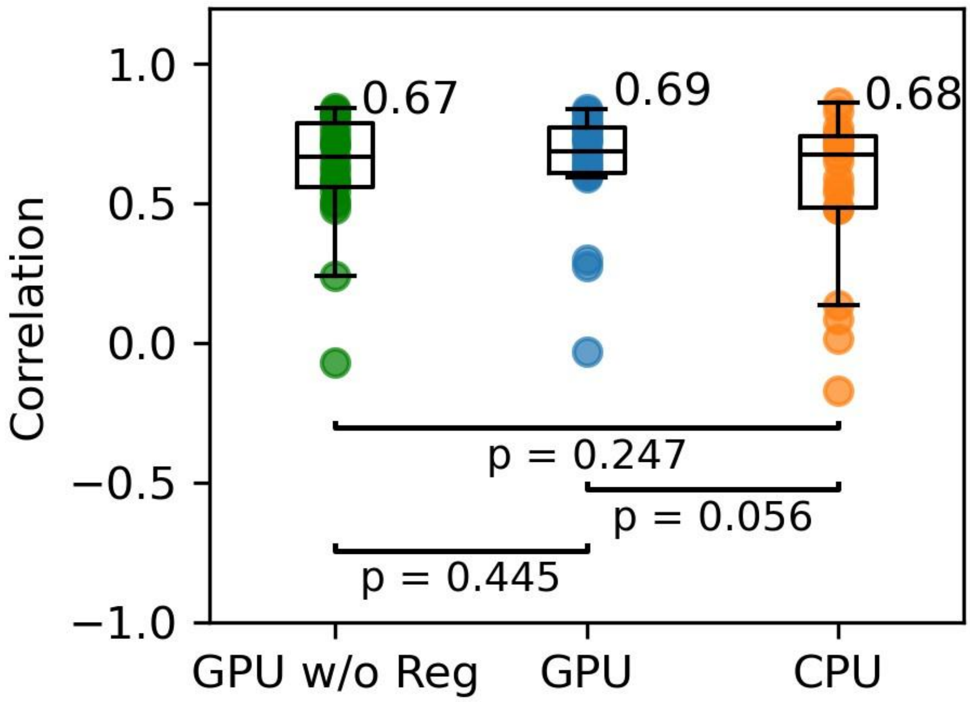


**Supplementary Figure S5. Effects of removing regularization on performance** After having discovered that the new GPU implementation allowed us to remove the steady state and state regulation effects, we decided to see how removing these affected performance. The overall effect gave a less powerful model than the model with the loss terms included but the effects were minor.

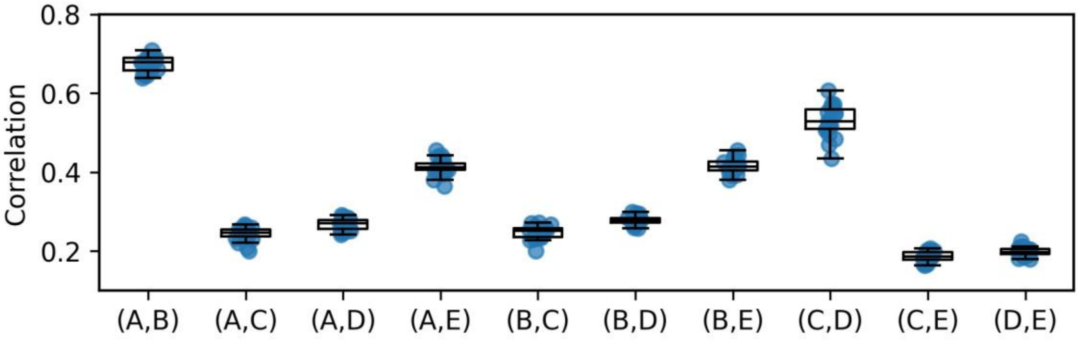


**Supplementary Figure S6. Comparison of internal states.** After training models using different conditions (A) the original version, (B) New state prior, (C), new weight regularization, (D) both new state prior and weight regularization (E) the new output bias with data without sigmoid transform, we observe the correlation between the internal states of the models. Measuring across conditions we see that C and D correlation with other models are consistent among the lowest; the correlation internally of C and D is however high. This indicates that removing the modified L2 loss with the new gradient noise term seems to have a relatively large effect on the network.

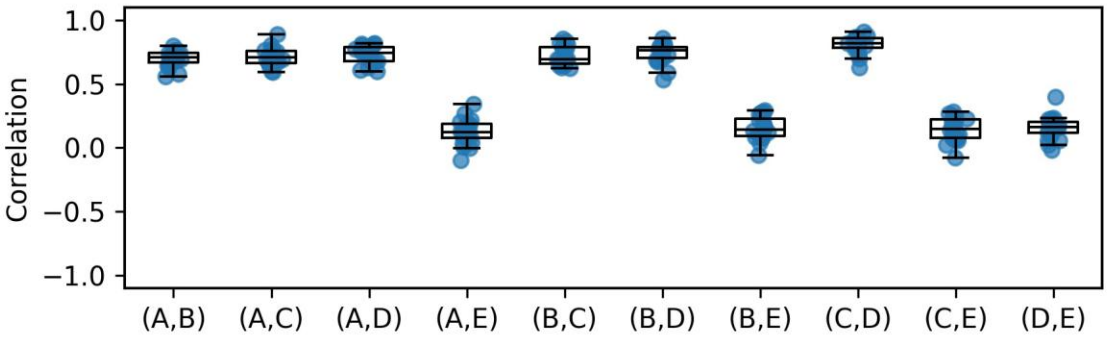


**Supplementary Figure S7. Comparison of output weights.** Above we show correlation between models outputs weights, these weights are used to scale TF activity as a final transform of the activities. The conditioned tested where (A) the original version, (B) New state prior, (C), new weight regularization, (D) both new state prior and weight regularization (E) the new output bias with data without sigmoid transform. Here we see that the introduction of the bias has a large effect on these weights. This should not be surprising as in the bias setting the data is rescaled.

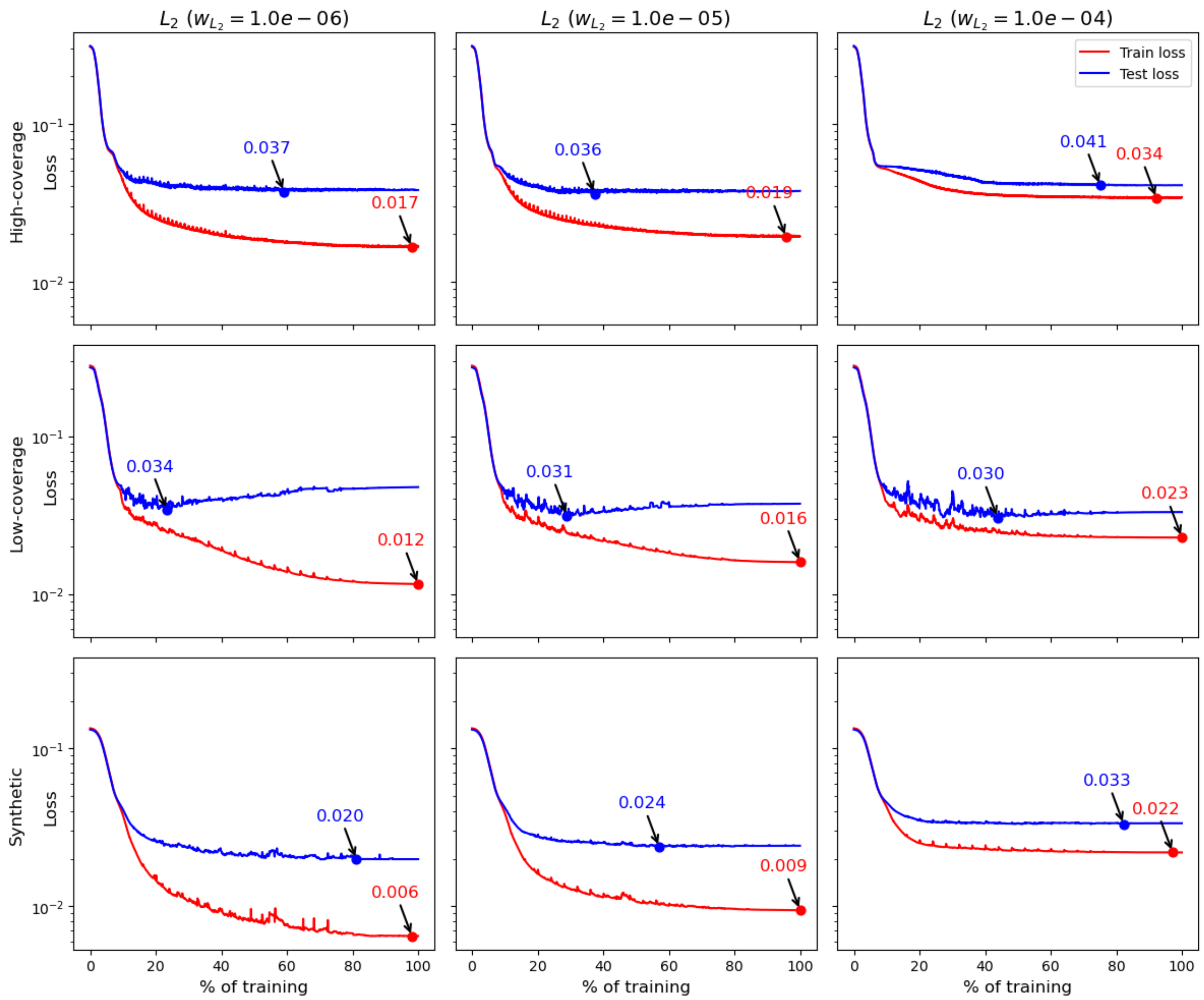


**Supplementary Figure S8. Loss curves for all regularization levels and datasets.** We suspected increasing the L2 loss causes a fall in the train loss, however the smallest test loss for different L2 norms differ between the datasets. Generally we see that a large effect on the L2 is first reached when we reach the largest L2 norm, here we also see that the smallest test loss recorded seem to be affected significantly in two out of the three cases.

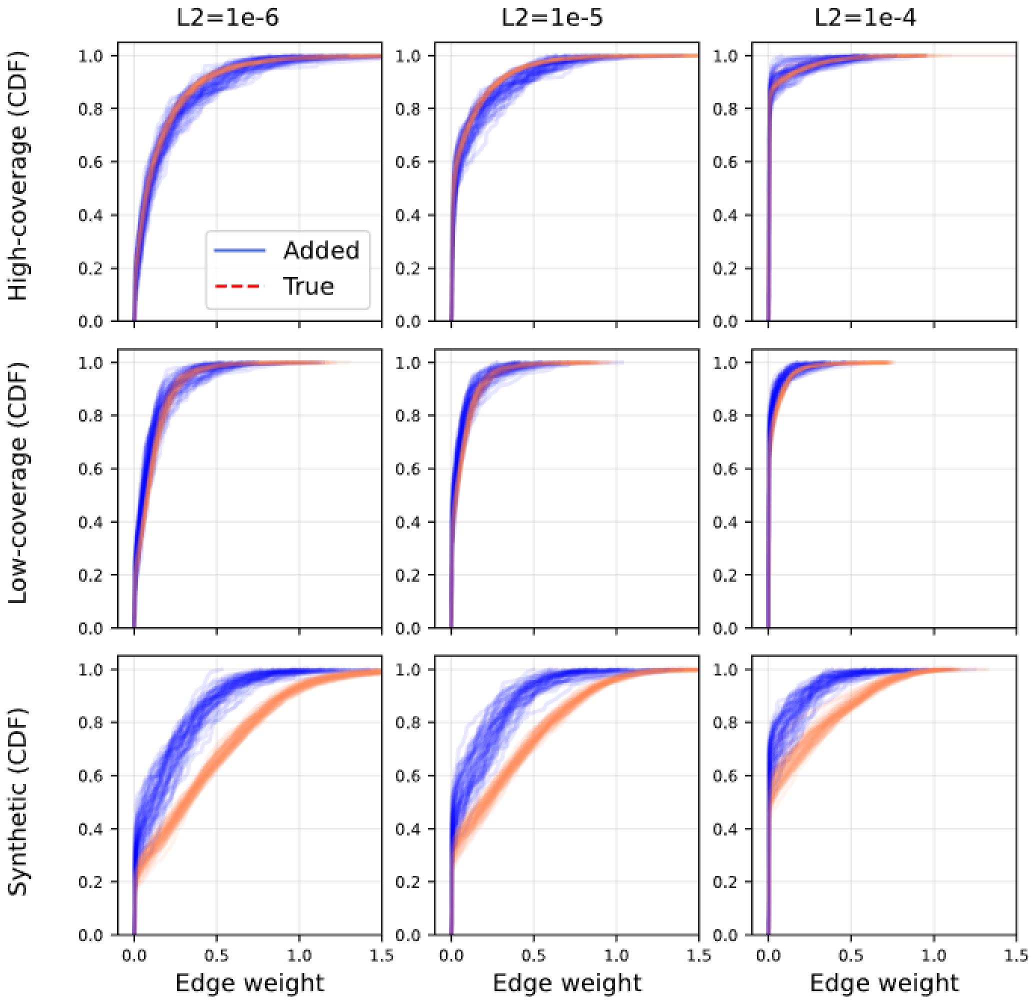


**Supplementary Figure S9. CDF for all regularization levels and datasets.** In the plot above we show the CDF for each model across all datasets and L2 settings.

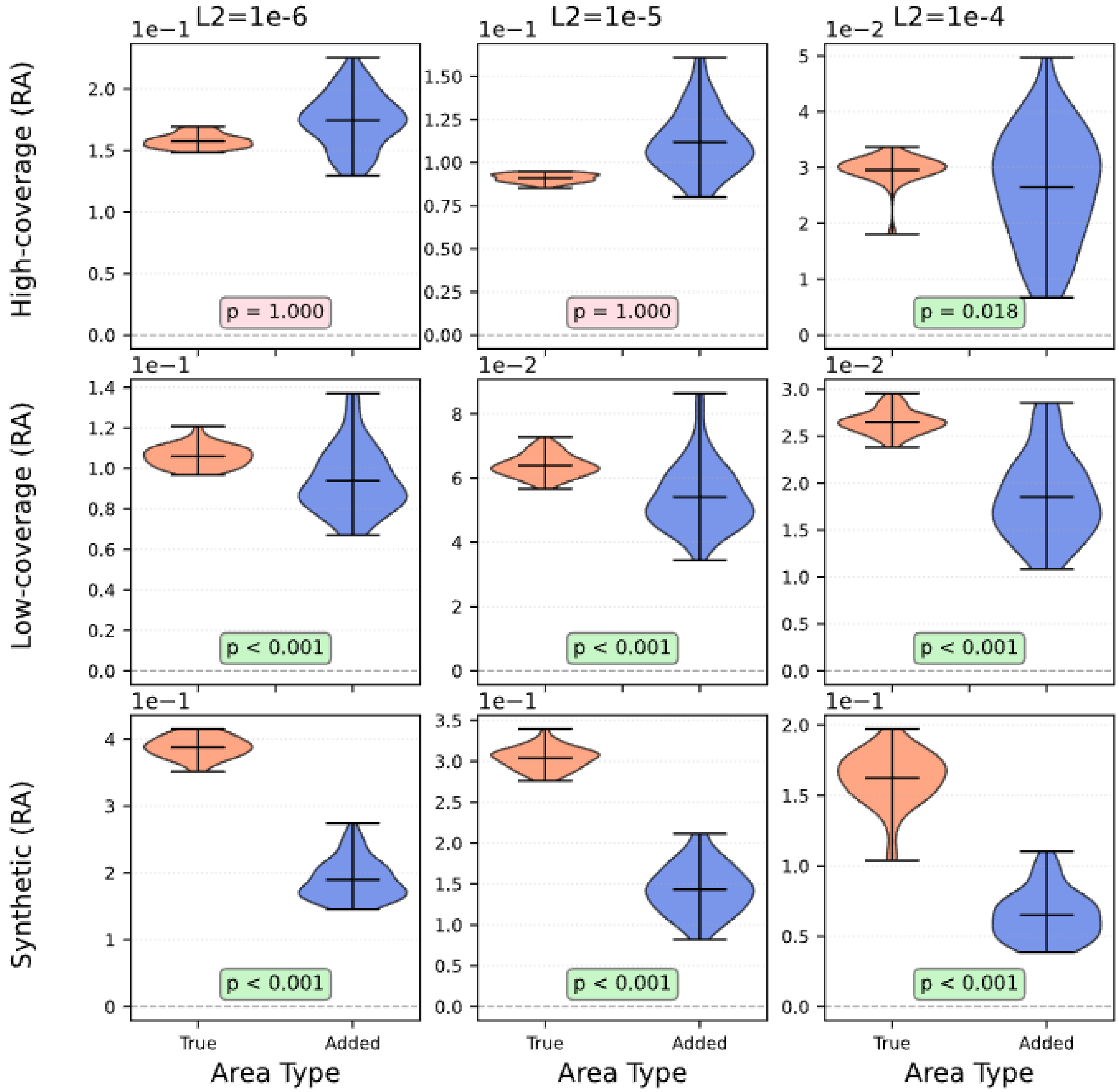


**Supplementary Figure S10. Violin for all regularization levels and datasets.** In the plot above we show the violin plot for each model across all datasets and L2 settings. The p value indicated is a one-sided Willcox test.

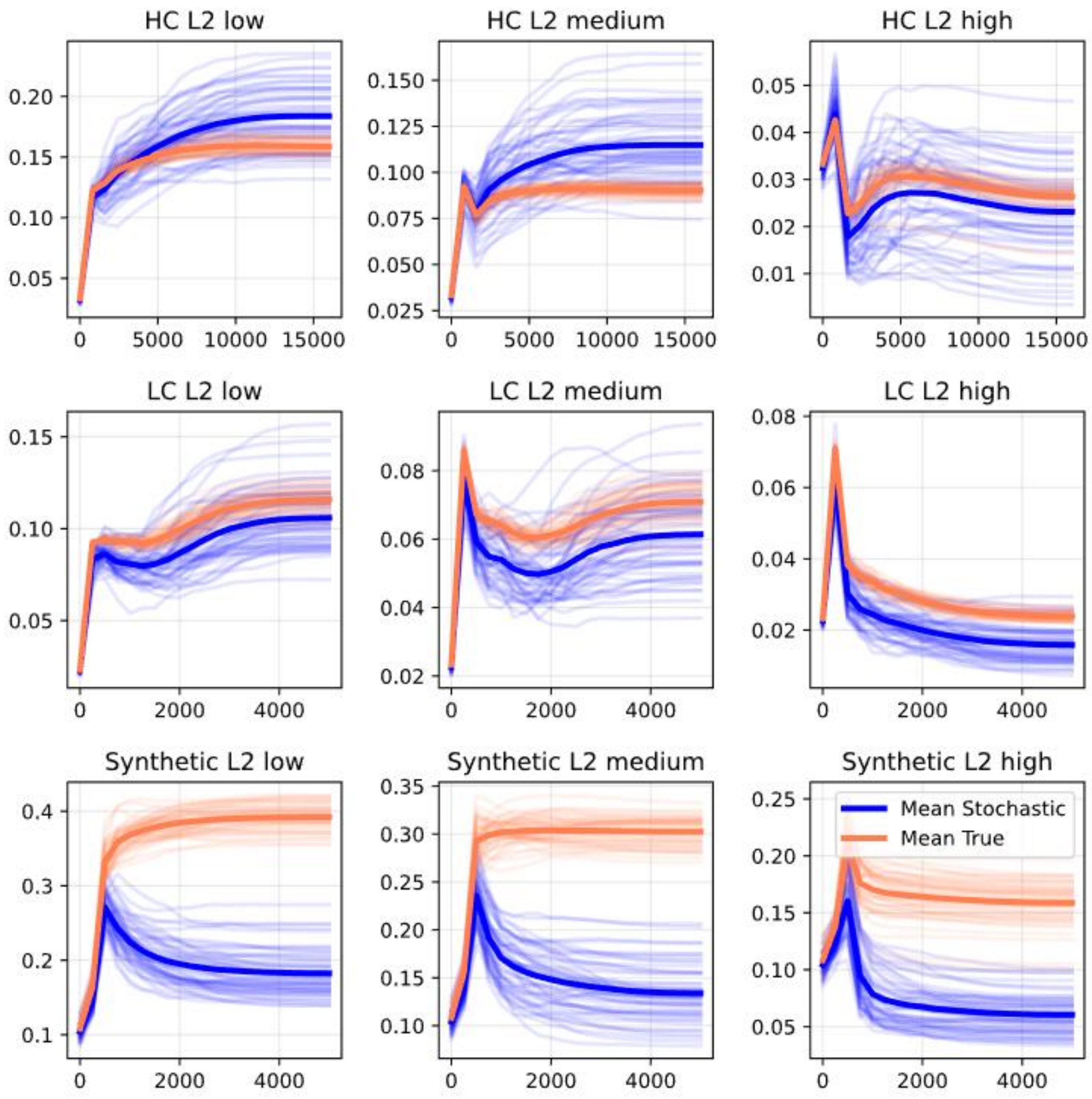


**Supplementary Figure S11. Residual Area over training for all regularization levels and datasets.** In the plot above we show the area over the checkpoints in training across all conditions we see that the plots start out next to each other and later diverse.

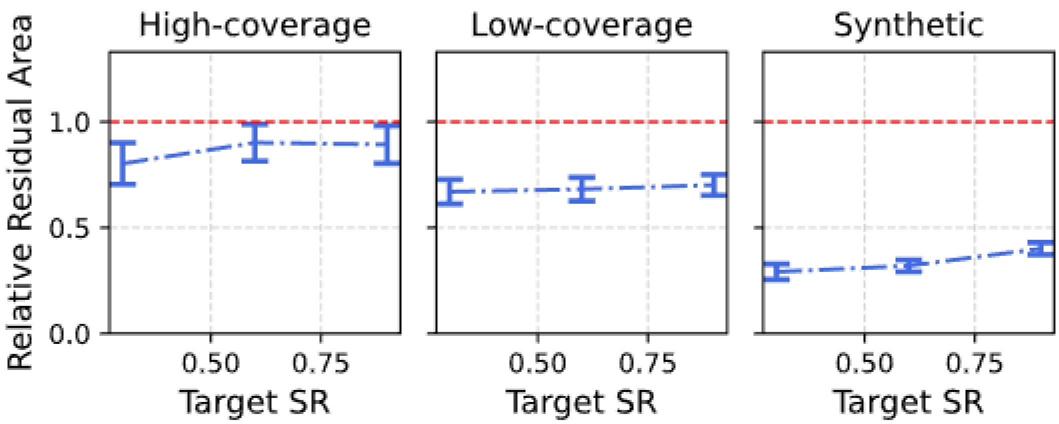


**Supplementary Figure S12. Relative Residual Area for varying target spectral radius.** In the plot we see that a smaller target for the spectral radius induces a smaller RRA. Having a smaller SR likely implies that there are weaker feedback mechanism in the model.

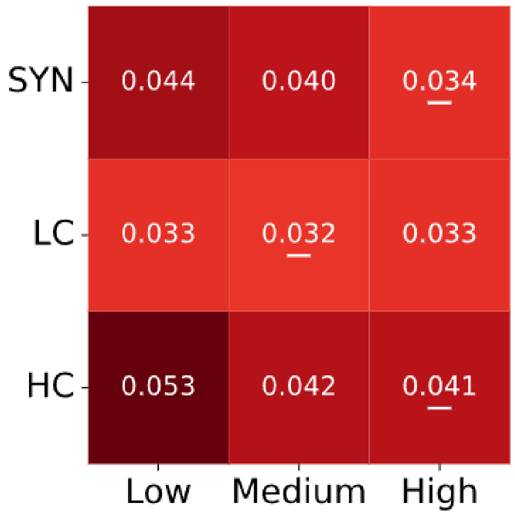


**Supplementary Figure S13. Best test loss for varying target spectral radius.** In the plot we show that best test loss for varying spectral radii (0.3,0.6 and 0.9). What this shows us is that there is a significant fall in performance for the synthetic and high coverage as the target spectral radius is decreased. This shows the limitation of lowering this hyperparameter that where similar the effect seen in the L2 study.

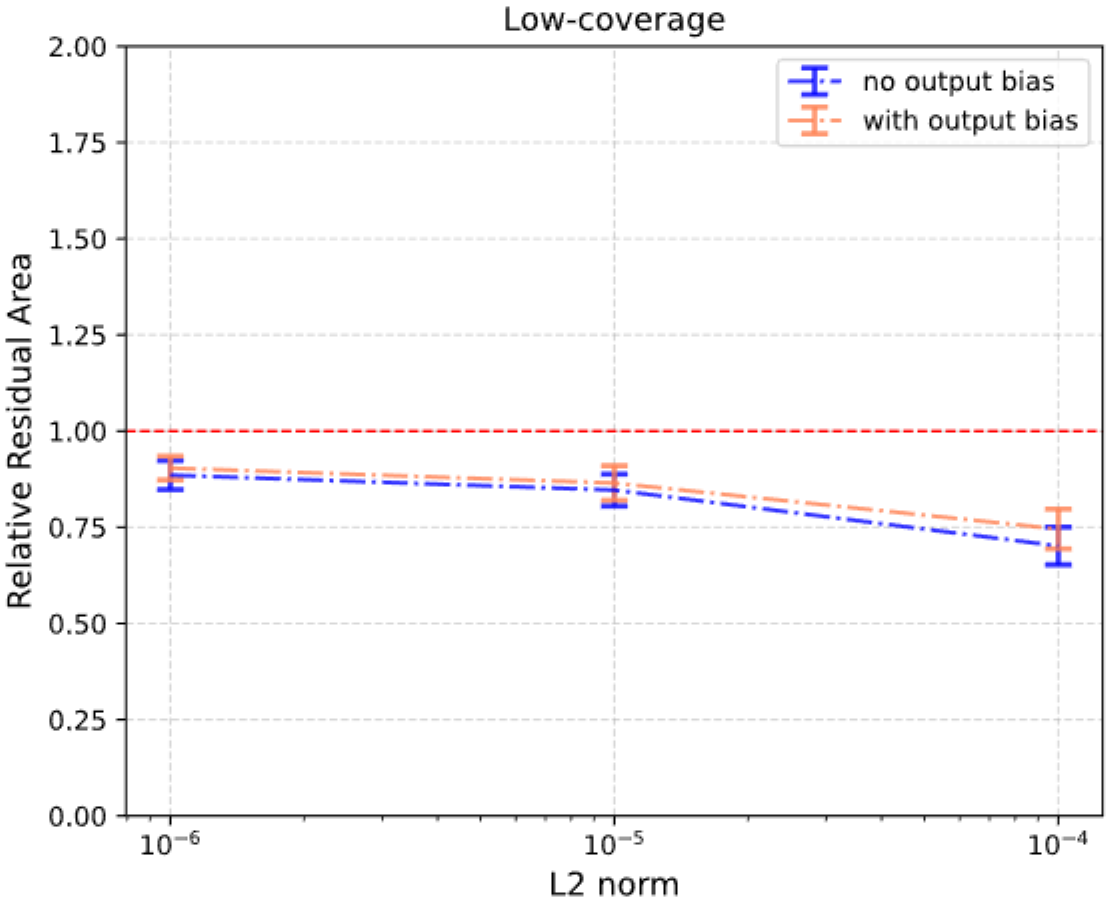


**Supplementary Figure S14. Comparing with and without the output bias using the RRA metric.** In the plot above we show that introducing the output bias seems to hurt performance in the low-coverage case however the effect is not significant as error bars overlap, error bars indicate $p<0.05$ region.

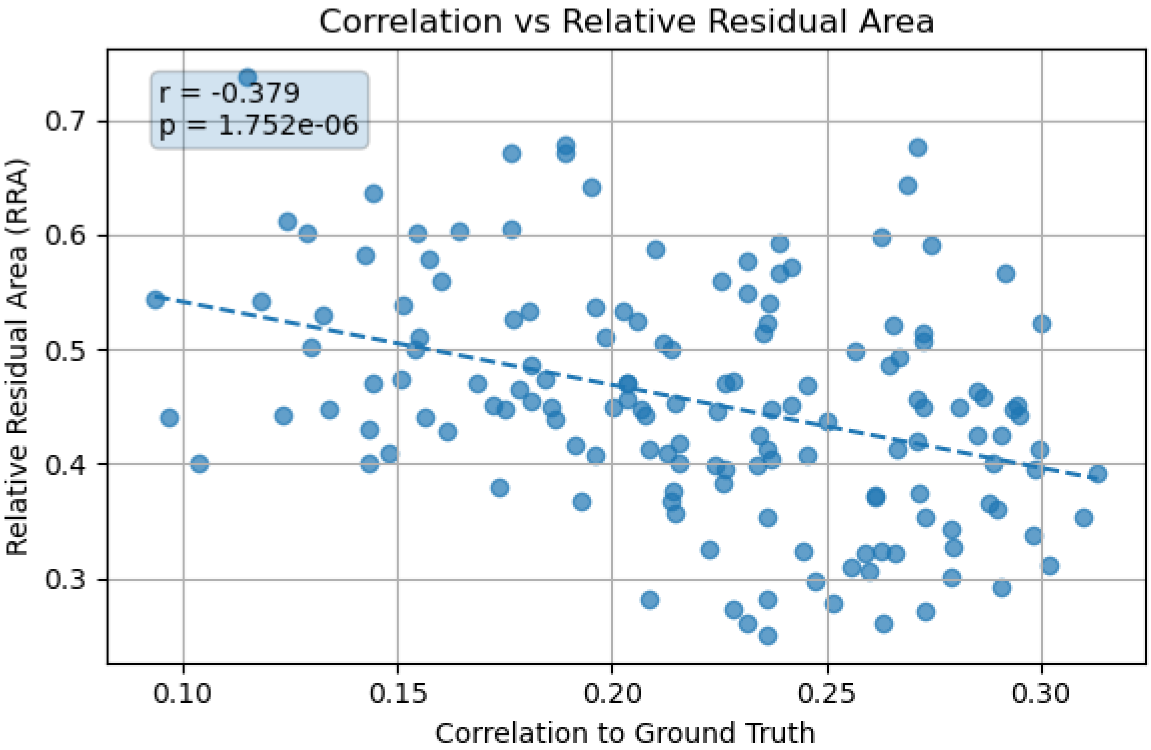


**Supplementary Figure S15. A lower RRA implies a larger correlation towards ground truth weights.** There is a negative correlation between the relative residual area and the correlation of the weights to the ground truth weights in the synthetically generated models. This negative correlation implies that if the RRA is low than there is a direct association with a more similar functional form to ground truth at least in the synthetic case.

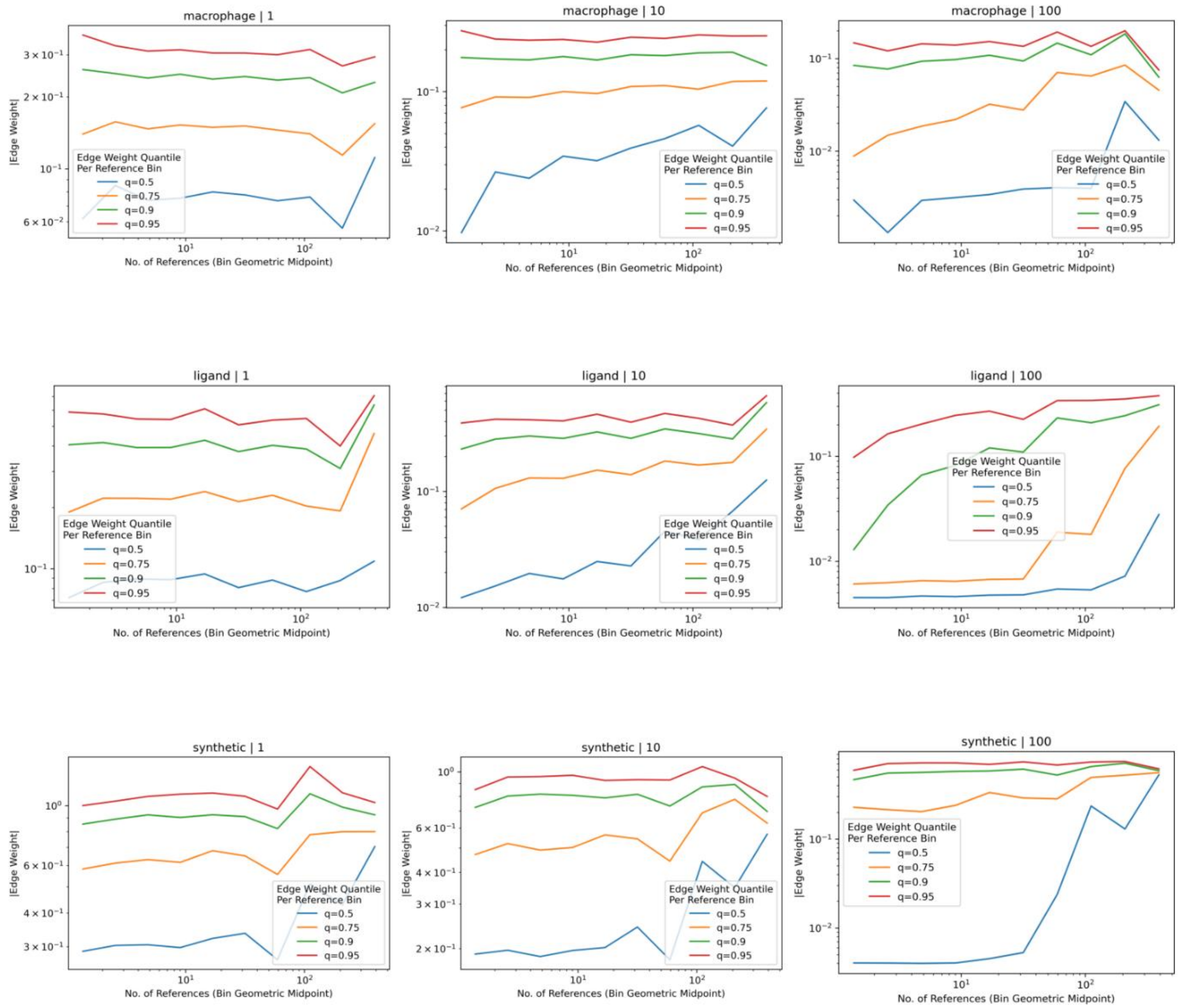


**Supplementary Figure S16. Relationship between prior literature support and learned edge weights.** Protein–protein interactions in the LEMBAS PKN were grouped into logarithmically spaced bins based on the number of literature references supporting each interaction. Within each reference bin, conditional quantiles of the absolute learned edge weight (q = 0.5, 0.75, 0.9, 0.95) were computed and plotted against the geometric midpoint of each bin on log–log axes. Lines show how different portions of the edge-weight distribution vary with increasing literature support across datasets.

**Algorithm 1** Conditioned Random Edge Addition

edge_list ← extract source, targets index from PKN
edge_MOA ← extract edge sign from PKN
**if** add_random = "conditioned" **then**
    *// Compute node degree weights*
    node_degrees ← bincount(edge_list.flatten(), minlength = $n$)
    node_weights ← $\frac{\text{node\_degrees}}{\sum \text{node\_degrees}}$
    *// Initialize existing edges set*
    existing_edges ← $\{(u, v) : (u, v) \in$ edge_list$\}$
    *// Generate new edges preferentially*
    new_edges ← [, ]
    **while** |new_edges| $< k$ **do** ▷ $k$ = edges_to_add
        $u \sim$ SampleInterger(node_weights) ▷ Sample source node
        $v \sim$ SampleInterger(node_weights) ▷ Sample target node
        **if** $u \neq v$ **and** $(u, v) \notin$ existing_edges **and** $(v, u) \notin$ existing_edges **then**
            new_edges ← new_edges $\cup \{(u, v)\}$
            existing_edges ← existing_edges $\cup \{(u, v)\}$
        **end if**
    **end while**
    *// Update edge list*
    edge_list ← edge_list ‖ new_edges$^T$ ▷ Horizontal concatenation
    *// Update edge MOA by sampling existing columns*
    indices $\sim$ Uniform$\{1, \ldots, |$edge_MOA$|\}^k$ ▷ Sample with replacement
    new_MOA ← edge_MOA[:, indices]
    edge_MOA ← edge_MOA ‖ new_MOA
**end if**

**Supplementary Algorithm A1 The pseudo code of edge addition.**

| | Parameter study - LC | Parameter study - HC | Self Pruning |
|---|---|---|---|
| State regularization | Included | Included | Removed |
| Gradient noise | dynamic scaled with learning rate | stable at 10**-2 | No edits |
| Batch size | 10 | 150 | No edits |
| Target Sprectral radius | 1.00E-05 | 1.00E-03 | No edits |
| Exp factor | 50 | 10 | No edits |
| Epochs trainied | 5000 | 16000 | No edits |
| Learning rate | 2*10**-3 | 1*10**-3 | No edits |
| Noise level | 10 | 1 | No edits |
| Uniform Lambda | 1.00E-05 | 1.00E-04 | No edits |
| Comment | These parameters where taken from the CPU study enterly | These parameters where first taken from the CPU implementation but later finetuned to accomiate a large batch size and still get similar performans to the CPU version. The process was to set the batch size to be maximally large and than change the parameter to better fit this large batch size. | The same parameters as in the previous two columns where used when No edits where made otherwise specfication are done |

**Supplementary Table 1 an overview of the experiment.**
